# Terminal Loop Sequences in Viral Double-Stranded RNAs Modulate RIG-I Signaling

**DOI:** 10.1101/2025.08.02.668285

**Authors:** Matthew Hackbart, Patrick Wang, Victoria Gnazzo, Carolina B. López

## Abstract

Detection of foreign RNAs is a crucial activation step for innate immunity pathways in response to viral infections. Retinoic acid-inducible gene I (RIG-I) is a cytoplasmic RNA sensor that triggers type I and III interferon (IFN) expression and activates the antiviral response in response to RNA virus infection. The activating ligand for RIG-I has been shown to be 5’-triphosphated, blunt-ended, double-stranded (ds)RNA, but questions remain on the impact of other RNA motifs on RIG-I activation. Here we show that immune-activating copy-back viral genomes (cbVGs) contain RNA stem loops away from the 5’ end of the RNA that enhance RIG-I signaling and IFN expression. Importantly, the sequence of the terminal loops of the activating motifs impacts the strength of IFN expression. Additionally, we show that synthetic versions of these cbVG-derived stem loops trigger innate immune responses in mice demonstrating their potential as immunostimulants *in vivo*.

## INTRODUCTION

Retinoic acid-inducible gene I (RIG-I) and melanoma differentiation-associated protein 5 (MDA5) are important cytoplasmic sensors of viral infections that initiate the antiviral immune response upon binding of viral RNA^1,2^. Binding of RIG-I or MDA5 by foreign RNA triggers a signaling cascade through the mitochondrial antiviral signaling (MAVS) protein that ends in the expression of antiviral proteins, including type I and III interferons (IFNs)^3–7^. RIG-I and MDA5 contain a C-terminal regulatory domain that recognizes the foreign RNA and drives conformational changes in the helicase domain to wrap around the RNA^8–13^. These conformational changes release caspase activation and recruitment domains (CARDs) that promote oligomerization of the proteins and signaling through MAVS^3,14,15^. RIG-I and MDA5 recognize unique pathogen-associated molecular patterns (PAMPs) found in foreign RNA but not in the host cell RNAs.

The most well-studied viral RNA PAMPs are within the RNAs that activate RIG-I. The regulatory domain of RIG-I detects terminal 5’ triphosphates on RNA^16–19^ and the detection of this motif can be inhibited by methylation of the 5’ end of the RNA^20,21^. While single-stranded (ss)RNA can activate RIG-I, the highest activation occurs when blunt-ended, double-stranded (ds)RNA^22^ contains the 5’ triphosphate^17^. The addition of overhangs to the end of the dsRNA limits activation of RIG-I^18,19^. RIG-I is activated best by short dsRNA while longer synthetic dsRNAs such as poly I:C (>500nt in length) activate MDA5^23–25^. With these guidelines in place, minimal ligands for activating RIG-I that contain a short 10 base pair dsRNA stem with a 5’ triphosphate have been synthetically generated^26,27^. While these studies have provided critical data to identify motifs that RIG-I recognizes and binds in the RNA, in general, they are not based on RNAs found naturally during viral infections. Further work is thus needed to determine whether other RNA structures or domains can further improve RIG-I recognition and identify the characteristics of viral RNAs that activate RIG-I signaling during actual infections.

Previously, we and others determined that the RNA that most robustly activates RIG-I signaling during Sendai virus (SeV) infection is a copy-back viral genome (cbVG)^28–30^. CbVGs contain complementary ends predicted to form a blunt-ended dsRNA PAMP with a 5’ triphosphate, characteristic of the canonical RIG-I ligand^31,32^. However, detailed folding analyses of the most prominent cbVG from SeV (cbVG 546) revealed a much more complex folding of the molecule, including an internal stem loop that is necessary for strong activation of the RIG-I pathway to stimulate IFN expression. This stem loop is comprised of nucleotides 70-114 of cbVG 546, and when transferred to inert RNAs, it significantly enhanced the immunostimulatory capability of the RNA^33^. Interestingly, the removal of stem loop 70-114 from cbVG 546 severely decreased RIG-I activation even though the cbVG still contained the 5’ triphosphate and predicted blunt-end dsRNA^33^.

It is unknown how SeV motif 70-114 enhanced RIG-I activity or whether other RIG-I stimulatory cbVGs contain stem loops similar to SeV 70-114. We therefore sought to determine if other viral cbVGs contain RNA stem loops that can transfer RIG-I stimulatory activity to otherwise inert RNAs and identify features of these RNA stem loops that are necessary for robust RIG-I activation. Here, we identified multiple sequences in cbVGs from human respiratory syncytial virus (RSV) and Nipah virus (NiV) infections that can transfer RIG-I stimulatory activity to the inert X RNA from the hepatitis C virus. Through mutations of the terminal loop sequences of the cbVG-derived RNAs, we discovered that specific nucleotides in the terminal loop of the RNA are necessary for strong activation of RIG-I signaling in cells. Lastly, *in vivo* administration of the cbVG-derived RNA loops activated innate immune signaling pathways, suggesting their potential use as immunostimulants for vaccines and immunotherapies.

## RESULTS

### RSV and NiV cbVGs stimulate interferon expression

To identify additional natural RNA stem loops that could activate antiviral innate immune pathways, we selected previously identified cbVGs from either RSV (RSV cbVG 236)^34^ and NiV (DI15, here called NiV cbVG 378)^35^. We first determined if these cbVGs can activate the IFN signaling pathway in human A549 lung cells and found that *in vitro* transcribed RNAs of both RSV cbVG 236 and NiV cbVG 378 could stimulate the transcription of type I (*IFNB1*) and type III (*IFNL1)* interferons at 6 hours post transfection. In contrast, the hepatitis C virus X region, a well characterized inert viral RNA^36,37^, did not stimulate IFN expression (Fig 1A and 1B). IFN expression was largely RIG-I and MAVS-dependent, similar to what we had reported for SeV cbVG 268, a shortened version of cbVG 546 (Fig 1C and 1D)^33^. The activation of this signaling pathway was also confirmed by the presence of phosphorylated TANK-binding kinase 1 (TBK1)(Fig 1E).

**Figure 1.**
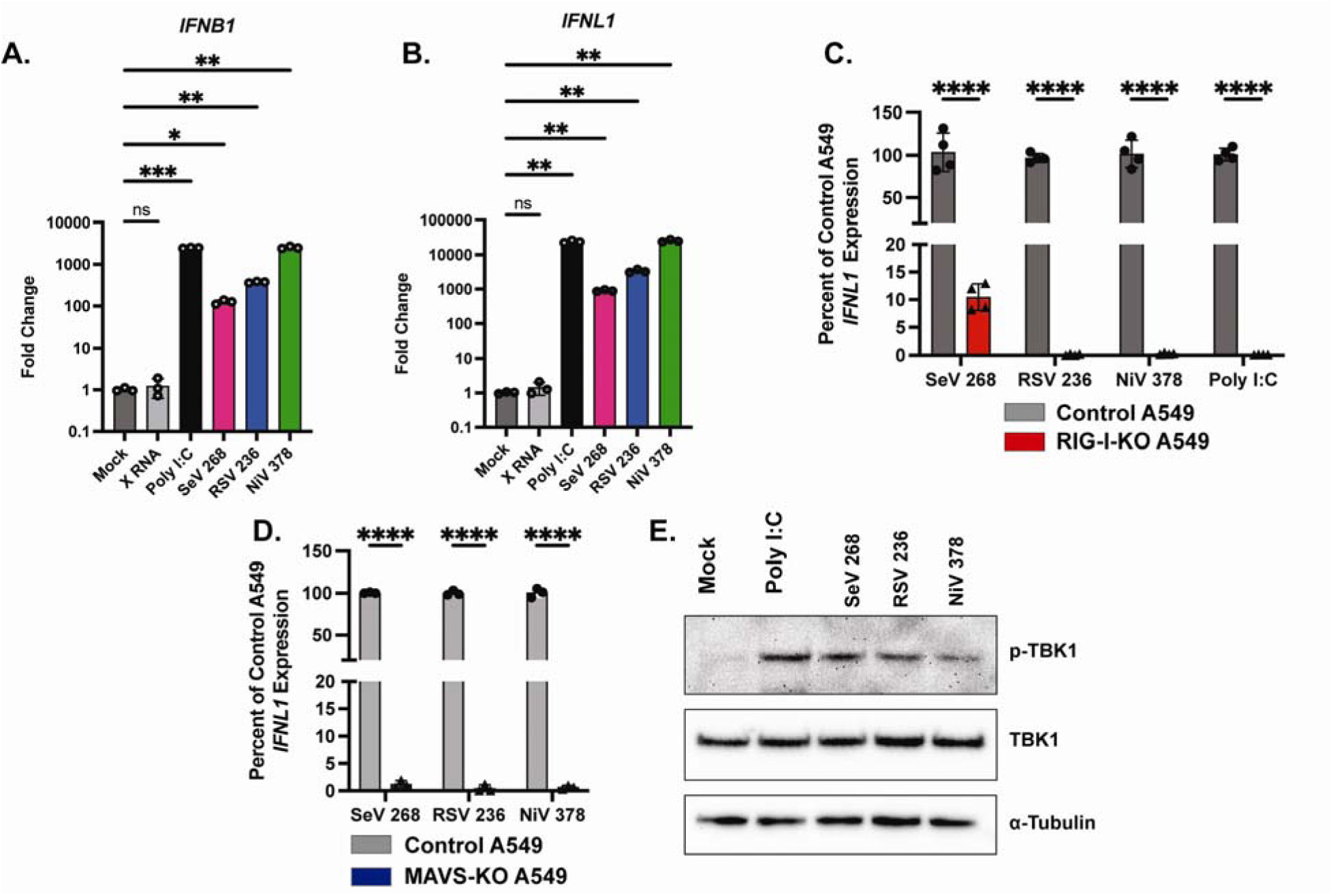
CbVG from RSV and NiV induce MAVS-dependent IFN expression. (A+B) Quantitative PCR (qPCR) for (A) type I interferon (*IFNB1*) and (B) type III interferon (*IFNL1*) expression by A549 cells at 6 hours post transfection (hpt) with 5 pmol of HCV X RNA, SeV cbVG 268, RSV cbVG 236, or NiV cbVG 378 or 100 ng of poly I:C. One-way ANOVA with Tukey’s HSD posthoc test was performed for statistical analysis. ns: p>0.05; *: p<0.05, **: p<0.01; ***: p<0.001. Data are represented as fold change over Mock. (C) qPCR for *IFNL1* of control or RIG-I-KO A549 cells at 6 hpt with 5 pmol of SeV cbVG 268, RSV cbVG 236, or NiV cbVG 378 or 100 ng of poly I:C. One-way ANOVA with Tukey’s HSD posthoc test was performed for statistical analysis. ****: p<0.0001. (D) qPCR for IFNL1 of control or MAVS-KO A549 cells at 6 hpt with 5 pmol of SeV cbVG 268, RSV cbVG 236, or NiV cbVG 378. One-way ANOVA with Tukey’s HSD posthoc test was performed for statistical analysis. ****: p<0.0001. Data reported are biological replicates. (E) Western blot cell lysates collected at 6hpt of A549 cells transfected with 5 pmol of SeV cbVG 268, RSV cbVG 236, or NiV cbVG 378 or 100 ng of poly I:C. Blots were stained for phosphorylated-TBK1, TBK1, and α-tubulin.

### RSV cbVGs contain immunostimulatory stem loops

To identify which specific RSV cbVG 236 sequence confers a strong RIG-I stimulatory activity to the molecule, we first made a series of deletions in cbVG 236 that either removed internal sequences or removed a complementary end of the cbVG (Fig 2A). We found that deleting the complementary ends of the cbVG (nucleotides 1-48 and 189-236) reduced RIG-I activation, as deletion mutants del 1-48 and 189-236 lost immunostimulatory activity (Fig 2B). Interestingly, we also found that nucleotides 49-100 are necessary for maximum IFN expression since deletions in this range (del 49-100, del 49-187, and del 64-187) showed reduced stimulation of the IFN response, but del 101-147 and del 137-187 induced IFN expression similar to the wild type cbVG 236.

**Figure 2.**
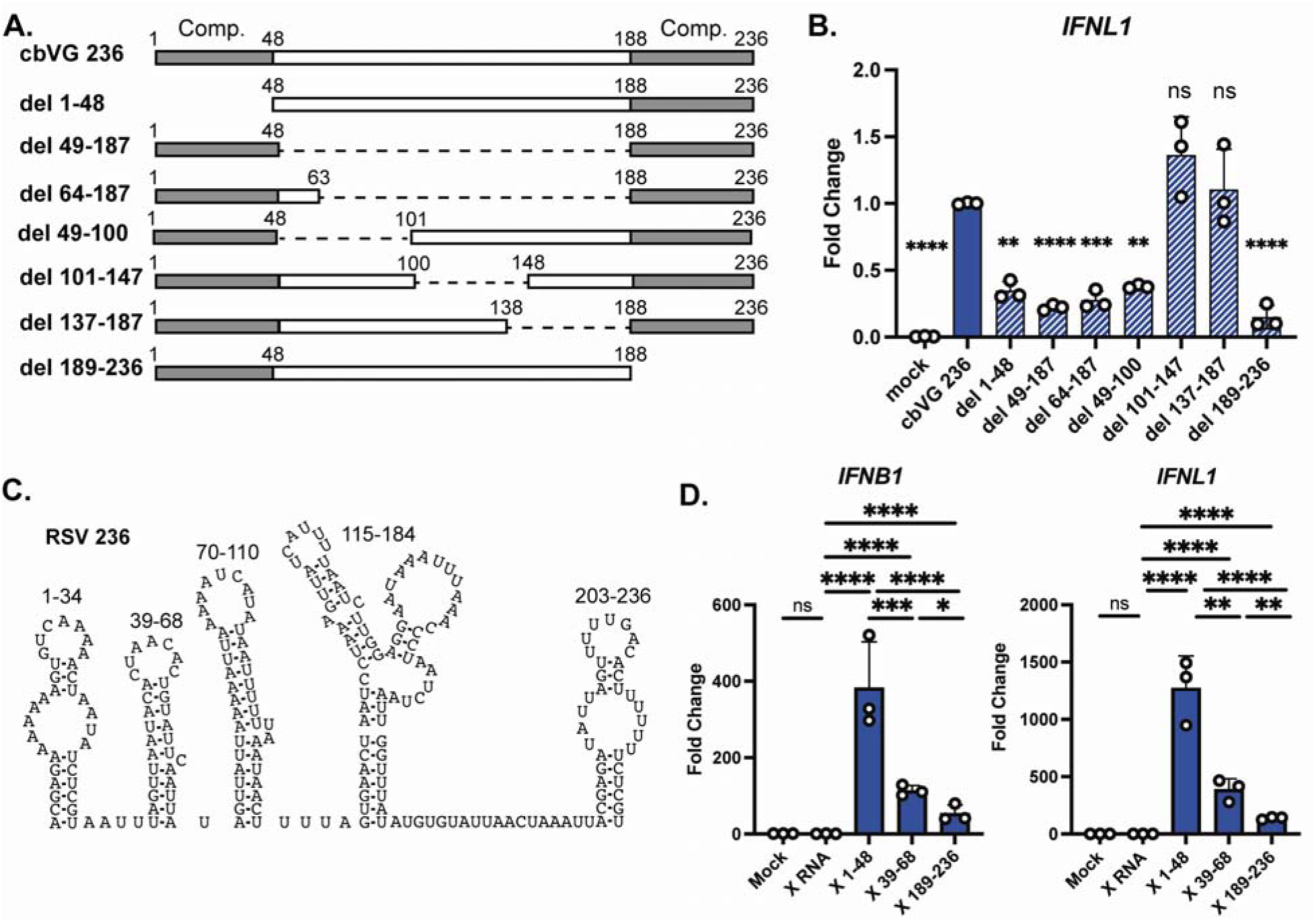
RSV cbVG sequences 1-48 and 39-68 are immunostimulatory. (A) Indicated deletions of RSV cbVG 236 were generated by *in vitro* transcription then 1 pmol of IVT RNA was transfected into A549 cells. Grey boxes represent complementary regions (Comp.) of the cbVG. (B) Expression of *IFNL1* was measured by qPCR at 6 hpt. One-way ANOVA with Tukey’s HSD posthoc test was performed for statistical analysis. ns: p>0.05; **: p<0.01; ***: p<0.001, ****: p<0.0001. Data are represented as fold change over cbVG 236. (C) Predicted structure of RNA stem loops from RSV cbVG 236. (D) Sequences from RSV cbVG 236 were attached to X RNA then *in vitro* transcribed. 1 pmol of RNA was transfected into A549 cells and expression of *IFNB1* and *IFNL1* was measured by qPCR at 6 hpt. One-way ANOVA with Tukey’s HSD posthoc test was performed for statistical analysis. ns: p>0.05; *: p<0.05; **: p<0.01; ***: p<0.001 ****: p<0.0001. Data are represented as fold change over the X RNA control. Data reported are biological triplicates.

To determine if specific immunostimulatory stem loops were present within the immunostimulatory sequences, we then performed *in silico* prediction of the RNA stem loops using RNAfold^38^. Since the complement ends of the cbVGs interfere with the *in silico* folding predictions based on minimal free energy, and it is likely that favorable secondary structures form during RNA synthesis rather than once the entire molecule has been produced, we determined all possible RNA structures on partial cbVGs lacking either the 3’ or 5’ complementary sequences. RSV cbVG 236 was predicted to form five RNA stem loops (Fig 2C and Fig S3) in the absence of 3’-5’ complementation with two of the stem loops (1-34 and 39-68) present within the immunostimulatory sequences. RNA folding of X region + RSV 1-48 predicted an immunostimulatory stem loop of sequence 1-34 similar to the full length cbVG (Fig S2).

To determine if the immunostimulatory sequences identified within cbVG 236 retained their activity outside the context of the original cbVG, we transferred sequences within the immunostimulatory regions containing stem loops, or control stem loop sequences from regions that showed no immunostimulatory activity, to the inert X RNA motif. We ensured that the overall structures of the final molecules were similar (Fig S2)^37^ and tested for their ability to induce IFN expression. We hypothesized that stem loops that were disrupted by the deletion of nucleotides 49-100 (such as stem loop 39-68) would induce the highest IFN expression. We found that X-region chimeras carrying RSV cbVG sequences 39-68 induced IFN expression when transfected into A549 cells, but the highest IFN signal was stimulated by transfer of nucleotides 1-48. Interestingly, the reverse complement of nucleotides 1-48 (nucleotides 189-236) induced weaker IFN expression in this context despite contributing to IFN induction in the context of the full cbVG, suggesting that the activity of this region depends on additional sequences present in the full length cbVG (Fig 2D). Consistent with data on full cbVGs, the X RNA constructs induced expression of *IFNL1* in a RIG-I dependent manner (Fig S1). Overall, these data suggest that while stem nucleotides 49-100 are crucial for IFN stimulation by cbVG 236, multiple unique RNA stem loops can activate an inert RNA to varying strengths of IFN activation.

### NiV cbVGs contain immunostimulatory stem loops

Similar to RSV, we made a series of deletions to remove specific RNA sequences from NiV cbVG 378 and determined which portions of the cbVG are necessary for strong IFN expression. We found that nucleotides within the complementary ends of the cbVG were necessary for strong expression of IFNs and removal of nucleotides 325-360 also decreased IFN expression (Fig 3A and B) suggesting that a stem loop in region 325-360 is necessary for a strong IFN response. NiV cbVG 378 is predicted to have eight RNA stem loops (Fig 3C and Fig S4). The deletion of region 325-360 is predicted to remove stem loop 324-361 from the cbVG, which suggests that this stem loop enhances IFN production during NiV cbVG 378 transfection. When attached to the X RNA, we again found that while NiV 325-360 induced *IFNB1* and *IFNL1* expression, other NiV sequences, such as 44-78 and 85-136 induced greater *IFNL1* expression (Fig 3D). Unexpectedly, 44-78 or 85-136 sequences, which are the strongest IFN-stimulating stem loops during X RNA attachment, were not necessary for IFN induction in the cbVG as deletion of either 44-78 or 81-140 did not reduce IFN expression. This suggests that either the RNA folds differently in the context of whole cbVG than when added X RNA stem loops, or other secondary structures impact the motifs activity. Further work is necessary to determine the exact structures of these RNAs in both contexts.

**Figure 3.**
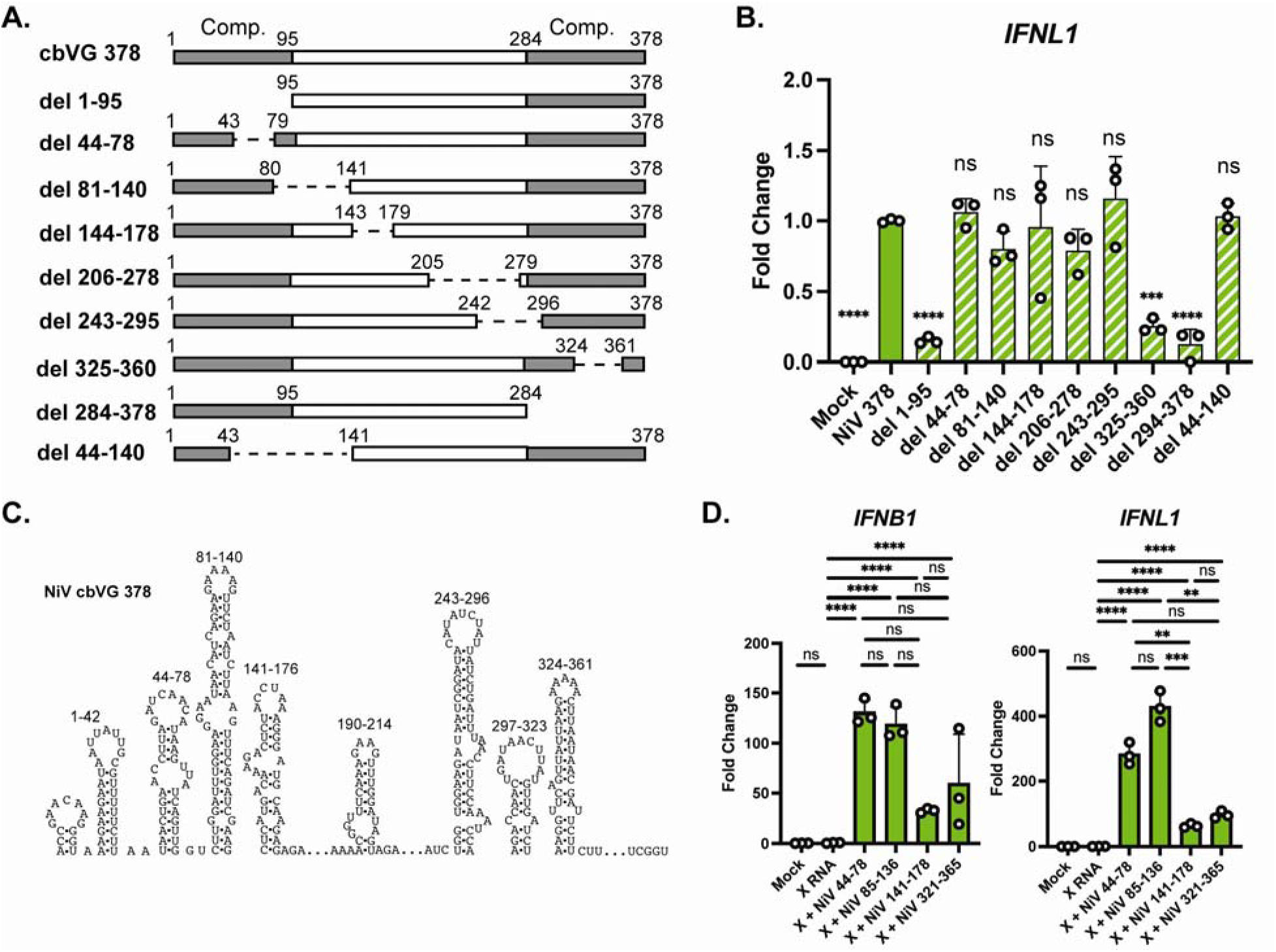
NiV cbVG sequences 44-78 and 85-136 are immunostimulatory. (A) Indicated deletions of NiV cbVG 378 were generated by *in vitro* transcription then 1 pmol of IVT RNA was transfected into A549 cells. Grey boxes represent complementary regions (Comp.) of the cbVG. (B) Expression of *IFNL1* was measured by qPCR at 6 hpt. One-way ANOVA with Tukey’s HSD posthoc test was performed for statistical analysis. ns: p>0.05; ***: p<0.001, ****: p<0.0001. Data are represented as fold change over NiV cbVG 378. (C) Predicted structure of RNA stem loops from NiV cbVG 378. (D) Sequences from NiV cbVG 378 were attached to X RNA then *in vitro* transcribed. 1 pmol of IVT RNA was transfected into A549 cells and expression of *IFNB1* and *IFNL1* was measured by qPCR at 6 hpt.. One-way ANOVA with Tukey’s HSD posthoc test was performed for statistical analysis. ns: p>0.05; **: p<0.01; ***: p<0.001, ****: p<0.0001. Data are represented as fold change over the X RNA control. Data reported are biological triplicates.

### Terminal loop sequences are critical for strong IFN expression

Since we observed multiple stem loops with different strengths of RIG-I activation, we next investigated the characteristics conserved among stronger immunostimulatory RNA sequences versus weaker immunostimulatory sequences (Fig 4A). The cbVG RNAs are predicted to form stem loops that are diverse in their size and shape. However, two complementary sequences that are predicted to form loops with the same structure (RSV 1-34 and RSV 203-236) exhibit different immunostimulatory capabilities (Fig 2C). We did not observe significant differences in the minimum free energy (MFE) between the stem loops, most of which had a predicted MFE between -6 and -10 kcal/mol. In addition, we did not detect increased levels of the stronger RNA stem loops by qPCR at 6 hours post transfection, suggesting that constructs with these stem loops do not have better transfection efficiency or increased stability in the cells (Fig S5). Upon observation, we found that the terminal loops with strong immunostimulatory activity contain either a CAA motif or poly-A stretch at the tip of the loop (Fig 4A). In contrast, the weaker immunostimulatory loops are instead U-rich at the terminal loop residues, except for NiV 324-361 which contains an A-rich terminal loop.

**Figure 4.**
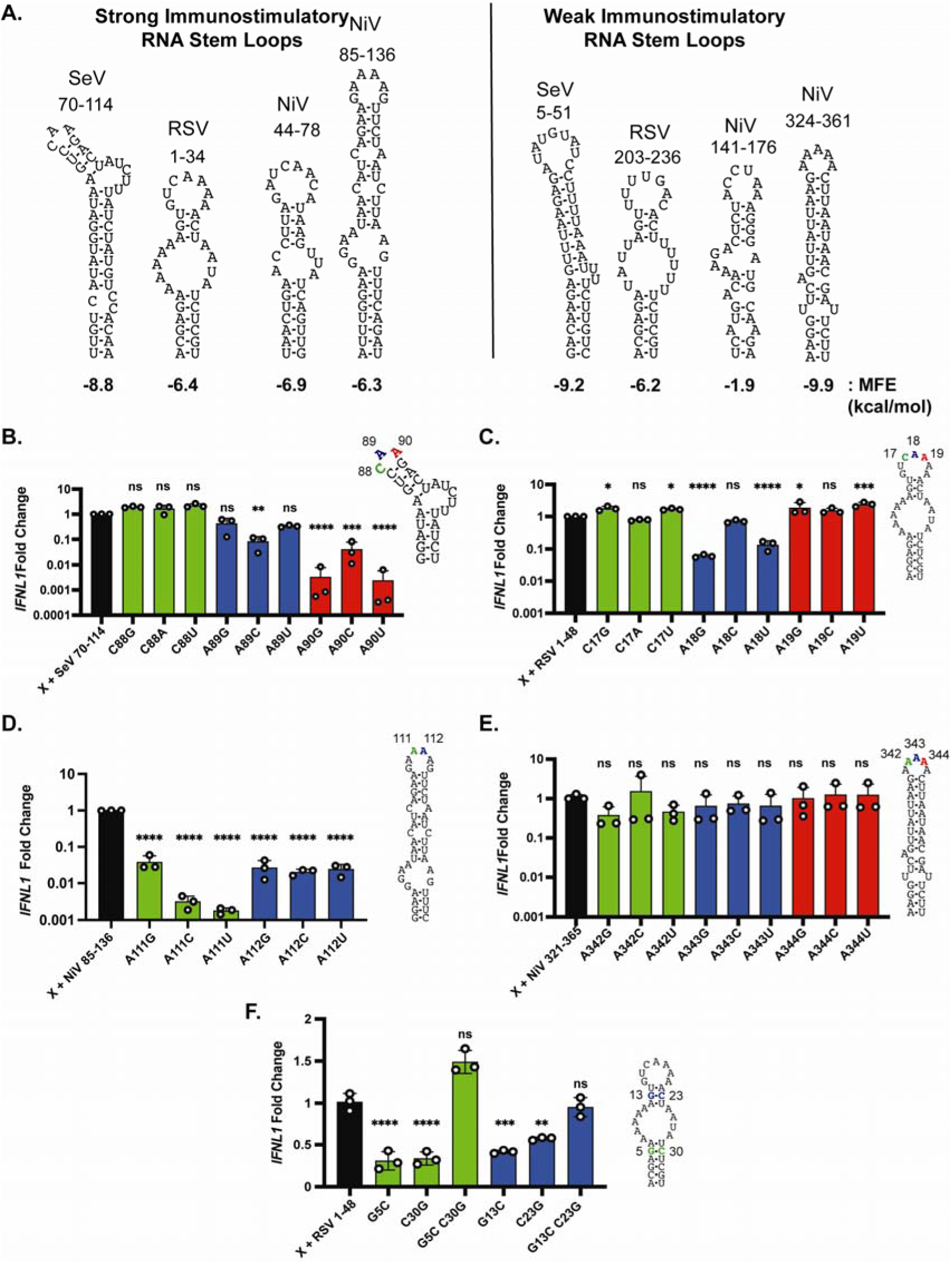
Terminal loop adenines are necessary for high IFN expression. (A) Predicted RNA structures of strong immunostimulatory and weak immunostimulatory stem loops derived from cbVGs. (B-F) Indicated mutations were generated for cbVG sequences attached to the X RNA. The loop sequences of (B) SeV 70-114, (C) RSV 1-48, (D) NiV 85-136, and (E) NiV 324-361 were mutated. (F) RSV 1-48 stem nucleotides were mutated to alter RNA structure. RNAs were *in vitro* transcribed and 1 pmol of IVT RNA was transfected into A549 cells. Expression of *IFNL1* was measured by qPCR at 6 hpt. One-way ANOVAs with Tukey’s HSD posthoc test were performed for statistical analysis. ns: p>0.05; *: p<0.05, **: p<0.01; ***: P<0.001; ****: p<0.0001. Data are represented as fold change over the wild-type stem loops. Data reported are biological triplicates.

We then hypothesized that the terminal loop sequences played a key role in triggering RIG-I signaling. To test this hypothesis, we mutated the terminal loops of the X-RNA attached sequences and tested their ability to induce IFN expression. Mutation of the terminal loop sequence were not predicted to alter the overall structure of the stem loops (Fig S6-9). For X + SeV 70-114, we found that mutation of the A89 or A90 terminal residues, specifically A89C, A90G, A90C, and A90U, greatly reduced IFN activation by the RNA construct, however mutation of the C88 residue did not (Fig 4B). Mutations in X + RSV 1-48 showed that A18G and A18U reduced IFN expression, but mutations of C17 or A19 slightly enhanced IFN expression (Fig 4C). Additionally, mutation of X + NiV 85-136 showed that both terminal adenines, A111 and A112, were necessary for IFN signaling (Fig 4D). However, the trend of terminal adenines was not observed for the weaker X + NiV 324-361, which did not show a reduction in IFN stimulation upon mutation of the terminal loop sequence (Fig 4E). These data suggest that the terminal loop sequences impact the strength of the IFN response.

Our previous work showed that the stems of the RNA structures were necessary for the SeV 70-114 stem loop to maintain strong IFN stimulation^33^. We therefore then determined if the stems were also necessary for X + RSV 1-48. We found that disruption of the stems by single mutations, G5C, C30G, G13C, and C23G, resulted in reduced IFN expression (Fig 4F) while complementing back the stem structure with double mutants, G5C C30G and G13C C23G, recovered IFN expression. These data suggest that the RNA structure as well as terminal loop sequence play a role a modulating IFN expression during RIG-I activation. Further work is necessary to determine if other features of these motifs are necessary for strong IFN induction.

### RSV 1-48 activates innate immune signaling *in vivo*

Previously, we found that the SeV cbVG 546 stem loop 70-114 activates a strong type I immune responses in mice when administered alone or within a lipid nanoparticle (LNP)^36,39,40^. Here, we aimed to determine whether RSV sequence 1-48 similarly activates innate sensors and induces an immune response *in vivo*. To ensure the RNA constructs were delivered intracellularly to trigger the RIG-I pathway, we packaged the X + RSV 1-48 and the control X + RSV 189-236 chimera constructs into LNPs (Fig 5A) and confirmed the homogeneity of the formulations before *in vivo* administration (Fig 5B). These formulations were then administered subcutaneously in the footpad of C57BL/6 mice. After 16 hours, we found that LNPs containing X + RSV 1-48 induced significantly higher expression of *IFNB1*, *MX1*, *IL1B*, and *CXCL10* compared to empty LNPs and X + RSV 189-236 (Fig 5C). These data demonstrate that the RSV cbVG-derived 1-48 sequence retains its immunostimulatory activity *in vivo* when packaged into LNPs and together with data from SeV 70-114 demonstrate that small RNA motifs with terminal loop adenines can serve as potent immunostimulators in mice.

**Figure 5.**
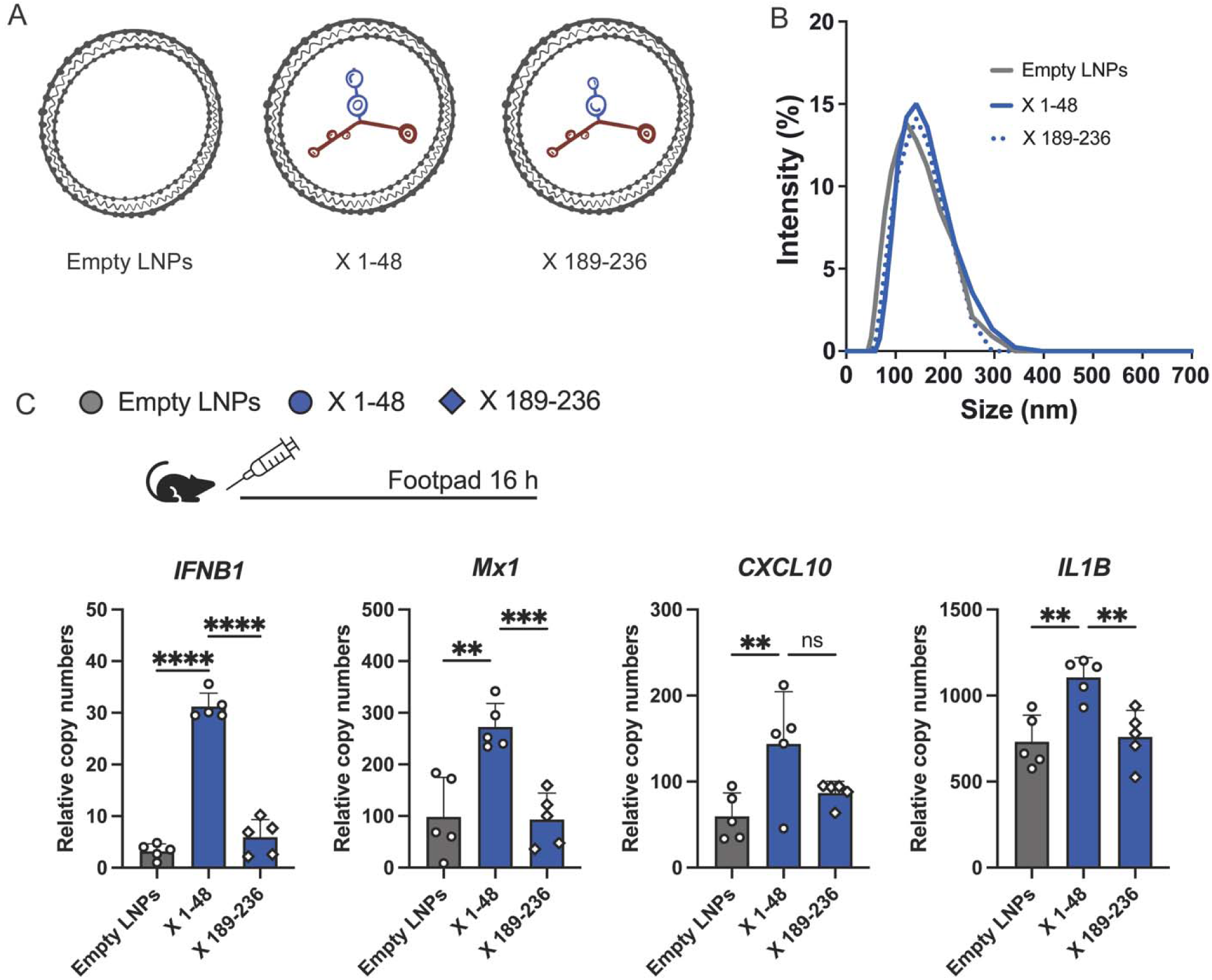
**X + RSV 1-48 activates innate immune signaling in mice. (**A) Schematic of lipid nanoparticles (LNPs) containing X + RSV 1-48 (X 1-48) or X + RSV 189-236 (X 189-236). (B) Dynamic light scattering measurement of LNP sizes of empty LNPs, X 1-48, and X 189-236. (C) LNPs were injected subcutaneously into the footpad of mice. Sixteen hours post injection, RNA was collected and the RNA expression of *IFNB1*, *MX1*, *CXCL10*, and *IL1B* was quantified by qPCR. One-way ANOVAs with Tukey’s HSD posthoc test were performed for statistical analysis. ns: p>0.05; **: p<0.01; ***: P<0.001; ****: p<0.0001. Data reported are 5 individual mice.

## DISCUSSION

We have identified multiple RNA stem loops derived from cbVGs of RSV and NiV that can transfer robust IFN stimulatory capabilities to an inert RNA. One unique feature of the higher immunostimulatory RNA stem loops is a richness for adenines at the tip of the loop. When these sequences were mutated, the mutant RNAs stimulated significantly less IFN upon transfection into cells. Upon packaging into lipid nanoparticles, the RSV cbVG-derived stem loop was able to activate innate immune signaling in mice.

While the enhanced stimulation appears to occur with adenine-rich sequences, we also observe that a stem loop with 5 adenines (NiV 325-360) did not have decreased IFN activation when the adenines were mutated (Fig. 4E). Since an adenine-rich loop does not always enhance IFN expression, more work is necessary to determine the exact motif or motifs that make up an optimal terminal loop PAMP and the impact of distinct tertiary conformations. Some possibilities are that sequences in the stems of the RNA interact with the loop to alter binding or that host proteins are recognizing a specific RNA motif that is more than adenines.

One major question that remains is what step of the RIG-I pathway is impacted by these terminal adenine-rich sequence. We propose the following possibilities for how the terminal loop sequences may alter IFN expression: 1) RIG-I is heavily modulated by phosphorylation and ubiquitination by host proteins^41^, so the terminal loop may recruit or inhibit additional host factors from interacting and altering RIG-I function in cells. 2) MAVS has recently been shown to interact with host RNAs^42^. The terminal sequence may stabilize RIG-I binding to MAVS through RNA/MAVS interactions and enhance downstream signaling to boost IFN expression. 3) The terminal loop adenines may be prone to RNA modifications such as N6 methylation to generate N6 methyladenosine (m6A)^43^. m6A RNA modification can alter the stability of the RNA in a cell and modulate the binding of RNA binding proteins. 4) Other unknown host RNA-binding proteins support robust RIG-I signaling. Further work is necessary to parse out what interactions occur with this terminal loop sequence and why adenine-rich sequences enhance IFN expression from the RIG-I pathway.

With the advent of mRNA vaccines and the usage of synthetic RNAs as therapeutics, understanding how RNAs can activate innate immunity is crucial to optimizing RNA-based adjuvants and treatments. Synthetic RNAs can act as immune adjuvants during vaccination^27,36,39,40,44,45^, and the addition of immunostimulatory stem loops to mRNA vaccines may bypass the need for additional adjuvants such as alum. Although additional work is necessary to assess the immunostimulatory properties of RNA stem loops *in vivo*, our discovery that the terminal loop sequence modulates immune activation can lead to the development of strategies to tailor the immune response not only based on dosage of stimulatory RNA, but on the sequence of attached RNA stem loops.

## MATERIALS AND METHODS

### Cells and Reagents

A549 human type II alveolar cells (ATCC, CCL-185), RIG-I-KO A549 cells (gifted from S. Weiss), and MAVS-KO A549 cells (gifted from S. Weiss) were cultured at 37°C and 5% CO_2_ with Dulbecco’s modified Eagle’s media (Thermofisher, 11995065) supplemented with 10% fetal bovine serum (FBS), 1 mM sodium pyruvate, 2 mM L-Glutamine, and 50 ug/mL gentamicin. All cell lines were treated with mycoplasma removal agent (MP Biomedicals, 093050044) and routinely tested for mycoplasma before use. Antibodies used in this study are as follows: rabbit anti-p-TBK1 (Cell Signaling, 5483, 1:2,000), rabbit anti-TBK1 (Abcam, AB40676, 1:2,000), rabbit anti-αtubulin (Invitrogen, 13-8000, 1:2,000). and goat anti-rabbit IgG-HRP (Cell Signaling, 7074, 1:10,000).

### *In Vitro* Transcription of RNA

RNAs were cloned into a pSL1180-T7 plasmid that contains T7 promoter at the 5’ end of the RNA sequence and hepatitis D virus ribozyme after the RNA sequence. *In vitro* transcription was performed with the Megascript T7 Transcription Kit (Thermofisher, AM1334) and RNA was isolated with LiCl precipitation according to manufacturer’s protocols. Purified RNA was measured by Qubit and tested for quality by Bioanalyzer (Agilent). Low Molecular Weight Poly I:C (Invivogen, tlrl-picw) was used as positive control for RIG-I activation. Full protocol available Protocols.io: dx.doi.org/10.17504/protocols.io.3byl49qprgo5/v2

### RT-PCR and qPCR

A549 cells were transfected with 5 pmol of indicated RNA and Lipofectamine 2000 (Thermofisher, 11668027). After 6 hours, RNA was isolated with Trizol (Thermofisher, 15596026). cDNA was generated with random hexamers using the High-Capacity RNA to cDNA kit (Thermofisher, 4387406) according to the manufacturer’s protocols. For qPCR quantification, cDNA was quantified with Power SYBR Green Mix (Thermofisher, 4367660). *IFNL1, IFNB1, IL1B, MX1, and CXCL10* were normalized to a house keeping index calculated from *ACTB* and *GAPDH* genes. Sequences of primers are given in Table S1. Full protocol available at Protocols.io: dx.doi.org/10.17504/protocols.io.8epv5rxw6g1b/v1

### Mice

C57BL/6 mice (The Jackson Laboratory) were bred in house. All mice were sex and age matched. Both male and female mice were included in the experiments.

### Lipid Nanoparticle formulation

X + RSV 1-48 and X + RSV 189-236 were encapsulated in the GenVoy Ionizable Lipid Mix (ILM) using a NanoAssemblr Ignite machine (Precision Nanosystems) following the manufacturer’s instructions. Empty LNPs were performed and used as controls. Encapsulation efficiency and RNA concentration were tested by Ribogreen Assay (ThermoFisher). Final particle size was measure by Dynamic Light Scattering. Full protocol available at Protocols.io: dx.doi.org/10.17504/protocols.io.36wgqnn4kgk5/v1

### Mice immunization

Mice were anesthetized with isoflurane and injected subcutaneously (s.c.) into a rear footpad. For immune response studies mice were inoculated with 0.5 µg X 1-48; 0.5 µg X 189-236; Empty LNPs or PBS at a final volume of 30 µl per dose. RNA was extracted from footpad 16 hours after inoculation using TRIzol (Ambion Inc.).

### *In Silico* RNA Folding Prediction

All RNA folding predictions were performed using the University of Vienna RNAfold website with the ViennaRNA Package (Version 2.6.3). Structures shown are the minimum free energy structures. All sequences used for folding are given in Table S2.

Website: http://rna.tbi.univie.ac.at/cgi-bin/RNAWebSuite/RNAfold.cgi

### Statistical Analysis and Reproducibility

All statistics were calculated using GraphPad Prism. Version 9. Specific tests and significance values are indicated in each figure legend. Data reported are biological triplicates unless otherwise stated.

### Ethics statement

All described studies adhered to the Guide for the Care and Use of Laboratory Animal of the National Institute of Health. Institutional Animal Care and Use Committee, Washington University in St. Louis approved protocol 23-0083.

## Supporting information

All supplementary figures

## ACKNOWLEDGEMENTS

The authors acknowledge the use of the Chemical and Environmental Analysis Facility at Washington University in Saint Louis for instrumentation and analytical support.

## Funding

NIH T32 HL007317-44 (MH)

NIH T32 AI007163-45 (MH)

NIH A137062 (CBL)

WashU BJC Investigator Program (CBL)

## Notes

### Competing Interest Statement

The authors have declared no competing interest.

### Summary of Updates

New data was added to strengthen the conclusions

## BIBLIOGRAPHY

1 Kato, H., Takahasi, K. & Fujita, T. RIG-I-like receptors: cytoplasmic sensors for non-self RNA. Immunol Rev 243, 91–98 (2011). 10.1111/j.1600-065X.2011.01052.x

2 Thoresen, D. et al. The molecular mechanism of RIG-I activation and signaling. Immunol Rev 304, 154–168 (2021). 10.1111/imr.13022

3 Yoneyama, M. et al. The RNA helicase RIG-I has an essential function in double-stranded RNA-induced innate antiviral responses. Nat Immunol 5, 730–737 (2004). 10.1038/ni1087

4 Tanner, N. K. & Linder, P. DExD/H box RNA helicases: from generic motors to specific dissociation functions. Mol Cell 8, 251–262 (2001). 10.1016/s1097-2765(01)00329-x

5 Kawai, T. et al. IPS-1, an adaptor triggering RIG-I- and Mda5-mediated type I interferon induction. Nat Immunol 6, 981–988 (2005). 10.1038/ni1243

6 Seth, R. B., Sun, L., Ea, C. K. & Chen, Z. J. Identification and characterization of MAVS, a mitochondrial antiviral signaling protein that activates NF-kappaB and IRF 3. Cell 122, 669–682 (2005). 10.1016/j.cell.2005.08.012

7 Xu, L. G. et al. VISA is an adapter protein required for virus-triggered IFN-beta signaling. Mol Cell 19, 727–740 (2005). 10.1016/j.molcel.2005.08.014

8 Cui, S. et al. The C-terminal regulatory domain is the RNA 5’-triphosphate sensor of RIG-I. Mol Cell 29, 169–179 (2008). 10.1016/j.molcel.2007.10.032

9 Kowalinski, E. et al. Structural basis for the activation of innate immune pattern-recognition receptor RIG-I by viral RNA. Cell 147, 423–435 (2011). 10.1016/j.cell.2011.09.039

10 Wang, Y. et al. Structural and functional insights into 5’-ppp RNA pattern recognition by the innate immune receptor RIG-I. Nat Struct Mol Biol 17, 781–787 (2010). 10.1038/nsmb.1863

11 Lu, C., Ranjith-Kumar, C. T., Hao, L., Kao, C. C. & Li, P. Crystal structure of RIG-I C-terminal domain bound to blunt-ended double-strand RNA without 5’ triphosphate. Nucleic Acids Res 39, 1565–1575 (2011). 10.1093/nar/gkq974

12 Lu, C. et al. The structural basis of 5’ triphosphate double-stranded RNA recognition by RIG-I C-terminal domain. Structure 18, 1032–1043 (2010). 10.1016/j.str.2010.05.007

13 Luo, D. et al. Structural insights into RNA recognition by RIG-I. Cell 147, 409–422 (2011). 10.1016/j.cell.2011.09.023

14 Peisley, A., Wu, B., Yao, H., Walz, T. & Hur, S. RIG-I forms signaling-competent filaments in an ATP-dependent, ubiquitin-independent manner. Mol Cell 51, 573–583 (2013). 10.1016/j.molcel.2013.07.024

15 Ramanathan, A. et al. The autoinhibitory CARD2-Hel2i Interface of RIG-I governs RNA selection. Nucleic Acids Res 44, 896–909 (2016). 10.1093/nar/gkv1299

16 Kim, D. H. et al. Interferon induction by siRNAs and ssRNAs synthesized by phage polymerase. Nat Biotechnol 22, 321–325 (2004). 10.1038/nbt940

17 Pichlmair, A. et al. RIG-I-mediated antiviral responses to single-stranded RNA bearing 5’-phosphates. Science 314, 997–1001 (2006). 10.1126/science.1132998

18 Schlee, M. et al. Recognition of 5’ triphosphate by RIG-I helicase requires short blunt double-stranded RNA as contained in panhandle of negative-strand virus. Immunity 31, 25–34 (2009). 10.1016/j.immuni.2009.05.008

19 Schmidt, A. et al. 5’-triphosphate RNA requires base-paired structures to activate antiviral signaling via RIG-I. Proc Natl Acad Sci U S A 106, 12067–12072 (2009). 10.1073/pnas.0900971106

20 Devarkar, S. C. et al. Structural basis for m7G recognition and 2’-O-methyl discrimination in capped RNAs by the innate immune receptor RIG-I. Proc Natl Acad Sci U S A 113, 596–601 (2016). 10.1073/pnas.1515152113

21 Schuberth-Wagner, C. et al. A Conserved Histidine in the RNA Sensor RIG-I Controls Immune Tolerance to N1-2’O-Methylated Self RNA. Immunity 43, 41–51 (2015). 10.1016/j.immuni.2015.06.015

22 Marques, J. T. et al. A structural basis for discriminating between self and nonself double-stranded RNAs in mammalian cells. Nat Biotechnol 24, 559–565 (2006). 10.1038/nbt1205

23 Kato, H. et al. Length-dependent recognition of double-stranded ribonucleic acids by retinoic acid-inducible gene-I and melanoma differentiation-associated gene 5. J Exp Med 205, 1601–1610 (2008). 10.1084/jem.20080091

24 Kato, H. et al. Differential roles of MDA5 and RIG-I helicases in the recognition of RNA viruses. Nature 441, 101–105 (2006). 10.1038/nature04734

25 Binder, M. et al. Molecular mechanism of signal perception and integration by the innate immune sensor retinoic acid-inducible gene-I (RIG-I). J Biol Chem 286, 27278–27287 (2011). 10.1074/jbc.M111.256974

26 Kohlway, A., Luo, D., Rawling, D. C., Ding, S. C. & Pyle, A. M. Defining the functional determinants for RNA surveillance by RIG-I. EMBO Rep 14, 772–779 (2013). 10.1038/embor.2013.108

27 Linehan, M. M. et al. A minimal RNA ligand for potent RIG-I activation in living mice. Sci Adv 4, e1701854 (2018). 10.1126/sciadv.1701854

28 Baum, A., Sachidanandam, R. & García-Sastre, A. Preference of RIG-I for short viral RNA molecules in infected cells revealed by next-generation sequencing. Proc Natl Acad Sci U S A 107, 16303–16308 (2010). 10.1073/pnas.1005077107

29 Mercado-López, X. et al. Highly immunostimulatory RNA derived from a Sendai virus defective viral genome. Vaccine 31, 5713–5721 (2013). 10.1016/j.vaccine.2013.09.040

30 Sun, Y. et al. Immunostimulatory Defective Viral Genomes from Respiratory Syncytial Virus Promote a Strong Innate Antiviral Response during Infection in Mice and Humans. PLoS Pathog 11, e1005122 (2015). 10.1371/journal.ppat.1005122

31 Kolakofsky, D. Isolation and characterization of Sendai virus DI-RNAs. Cell 8, 547–555 (1976). 10.1016/0092-8674(76)90223-3

32 Lazzarini, R. A., Keene, J. D. & Schubert, M. The origins of defective interfering particles of the negative-strand RNA viruses. Cell 26, 145–154 (1981). 10.1016/0092-8674(81)90298-1

33 Xu, J. et al. Identification of a Natural Viral RNA Motif That Optimizes Sensing of Viral RNA by RIG-I. mBio 6, e01265–01215 (2015). 10.1128/mBio.01265-15

34 Sun, Y. et al. A specific sequence in the genome of respiratory syncytial virus regulates the generation of copy-back defective viral genomes. PLoS Pathog 15, e1007707 (2019). 10.1371/journal.ppat.1007707

35 Welch, S. R. et al. Inhibition of Nipah Virus by Defective Interfering Particles. J Infect Dis 221, S460–s470 (2020). 10.1093/infdis/jiz564

36 Fisher, D. G., Coppock, G. M. & López, C. B. Virus-derived immunostimulatory RNA induces type I IFN-dependent antibodies and T-cell responses during vaccination. Vaccine 36, 4039–4045 (2018). 10.1016/j.vaccine.2018.05.100

37 Stone, A. E. et al. Hepatitis C virus pathogen associated molecular pattern (PAMP) triggers production of lambda-interferons by human plasmacytoid dendritic cells. PLoS Pathog 9, e1003316 (2013). 10.1371/journal.ppat.1003316

38 Zuker, M. & Stiegler, P. Optimal computer folding of large RNA sequences using thermodynamics and auxiliary information. Nucleic Acids Res 9, 133–148 (1981). 10.1093/nar/9.1.133

39 Fisher, D. G., Gnazzo, V., Holthausen, D. J. & López, C. B. Non-standard viral genome-derived RNA activates TLR3 and type I IFN signaling to induce cDC1-dependent CD8+ T-cell responses during vaccination in mice. Vaccine 40, 7270–7279 (2022). 10.1016/j.vaccine.2022.10.052

40 Gnazzo, V. et al. DDO-adjuvanted influenza A virus nucleoprotein mRNA vaccine induces robust humoral and cellular type 1 immune responses and protects mice from challenge. mBio, e0358924 (2024). 10.1128/mbio.03589-24

41 Rehwinkel, J. & Gack, M. U. RIG-I-like receptors: their regulation and roles in RNA sensing. Nat Rev Immunol 20, 537–551 (2020). 10.1038/s41577-020-0288-3

42 Gokhale, N. S. et al. Cellular RNA interacts with MAVS to promote antiviral signaling. Science 386, eadl0429 (2024). 10.1126/science.adl0429

43 Boulias, K. & Greer, E. L. Biological roles of adenine methylation in RNA. Nat Rev Genet 24, 143–160 (2023). 10.1038/s41576-022-00534-0

44 Jiang, X. et al. Intratumoral delivery of RIG-I agonist SLR14 induces robust antitumor responses. J Exp Med 216, 2854–2868 (2019). 10.1084/jem.20190801

45 Mao, T. et al. A stem-loop RNA RIG-I agonist protects against acute and chronic SARS-CoV-2 infection in mice. J Exp Med 219 (2022). 10.1084/jem.20211818

