## Supplementary material for "Terminal Loop Sequences in Viral Double-Stranded RNAs Modulate RIG-I Signaling": All supplementary figures

**SUPPLEMENTAL MATERIAL**

**Table S1. qPCR Primer Sequences**

**Table S2. Sequences of RNA for RNAfold**

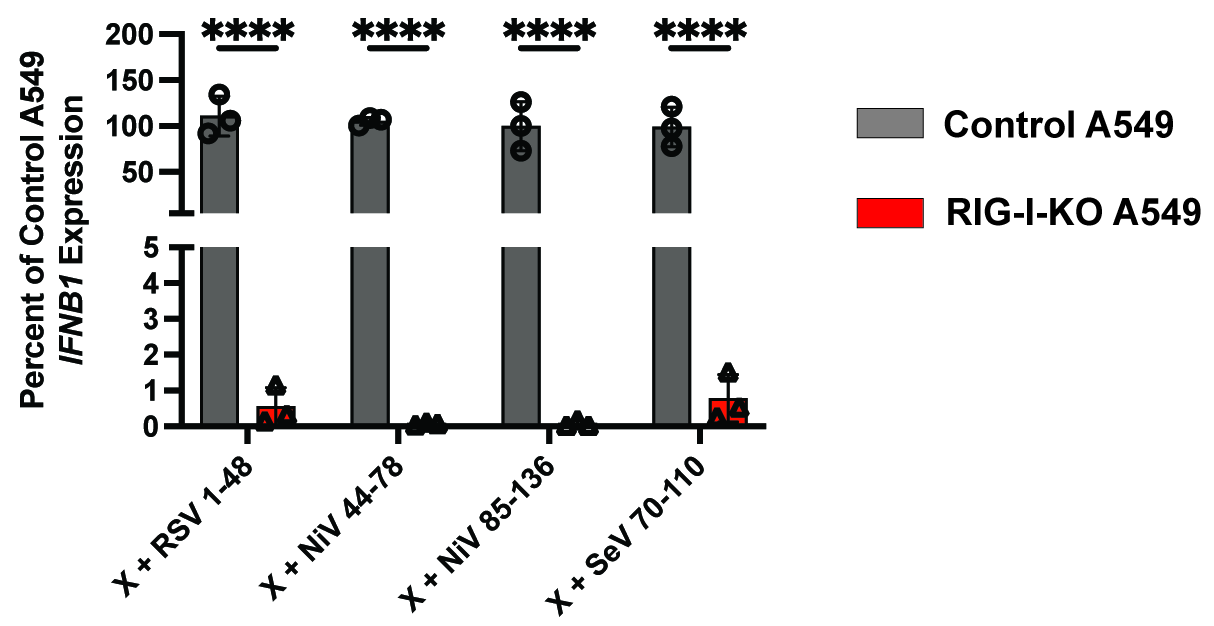

**Figure S1. X RNA constructs induces a RIG-I-dependent interferon response.** qPCR for IFNL1 of RNA collected from control or RIG-I-KO A549 cells at 6 hpt with 5 pmol of X RNA constructs. One-way ANOVA with Tukey’s HSD posthoc test was performed for statistical analysis. ****: p<0.0001. Data reported are biological triplicates.

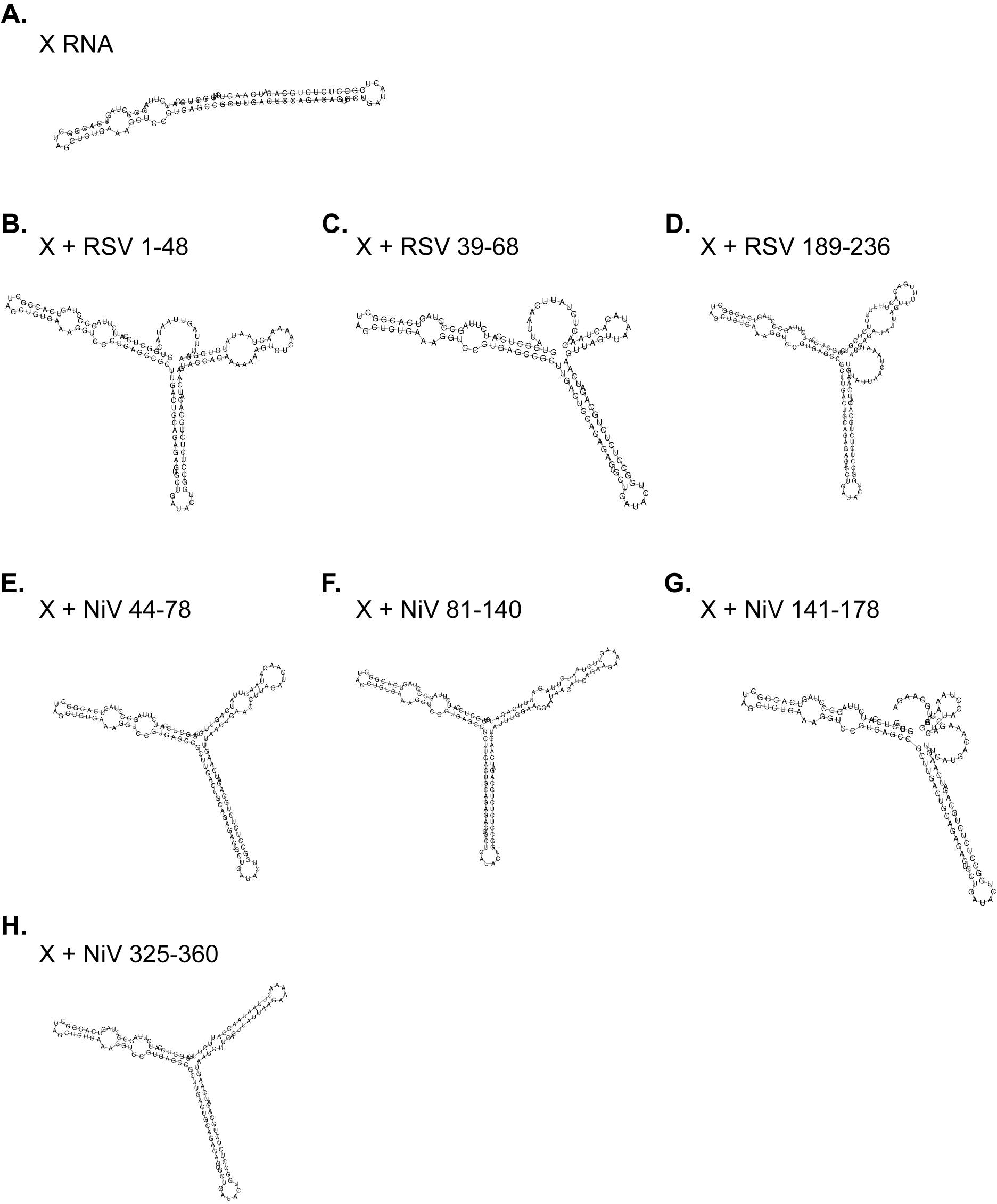

**Figure S2. Predicted RNA structures for X RNA-attached stem loops.** RNA fold predictions for (A) HCV X RNA, (B) X RNA with RSV stem loop 1-34, (C) X RNA with RSV stem loop 39-68, (D) X RNA with RSV stem loop 203-236, (E) X RNA with NiV stem loop 44-78, (F) X RNA with NiV stem loop 81-140, (G) X RNA with NiV stem loop 141-178, and (H) X RNA with NiV stem loop 325-360.

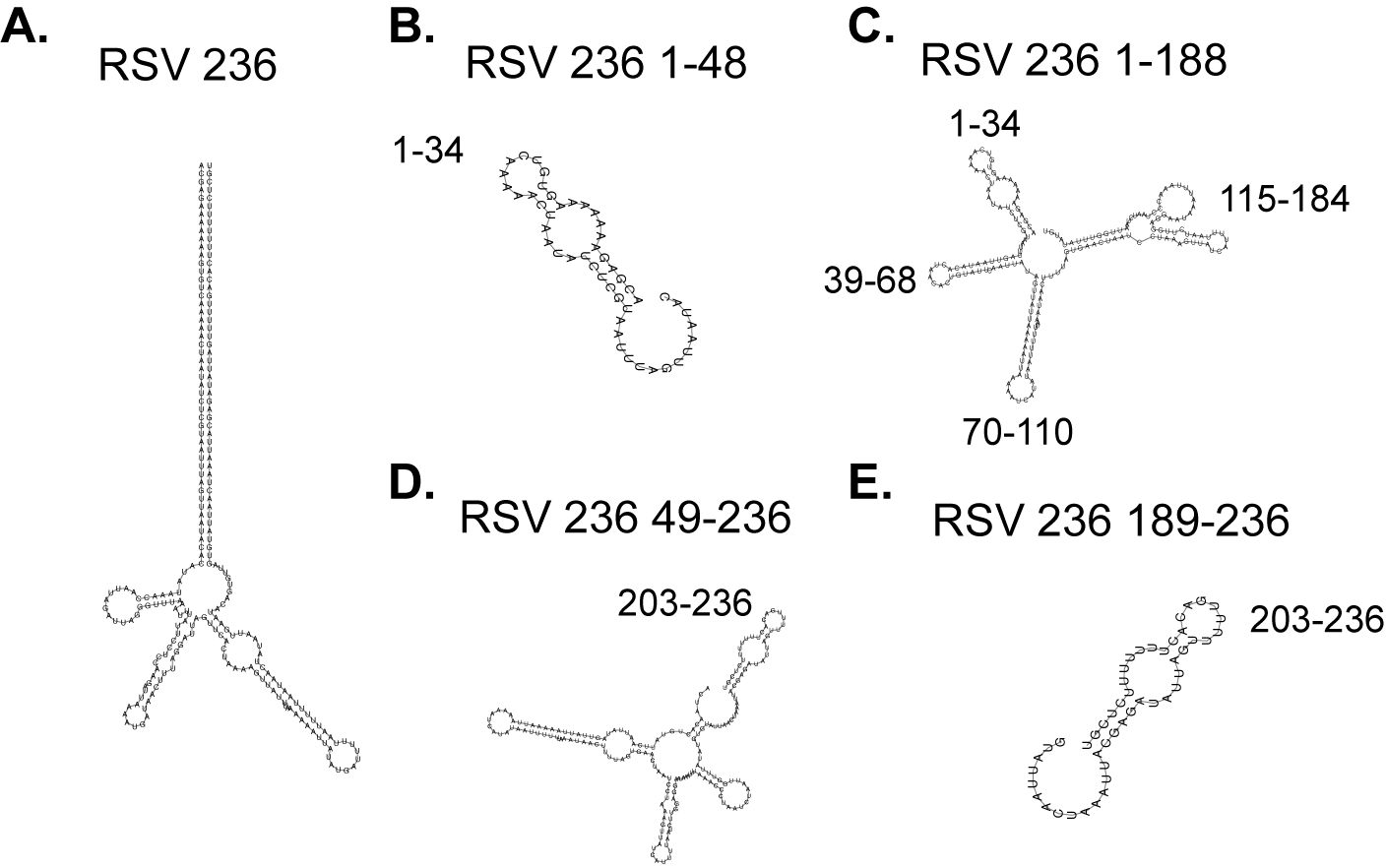

**Figure S3. Predicted RNA structures of RSV cbVG 236**. RNAfold predictions for the (A) full-length RSV cbVG 236, (B) nucleotides 1-48 of RSV 236, (C) nucleotides 1-188 of RSV 236, (D) nucleotides 49-236 of RSV 236, and (E) nucleotides 189-236 of RSV 236. Stem loops found in Fig 2A are indicated.

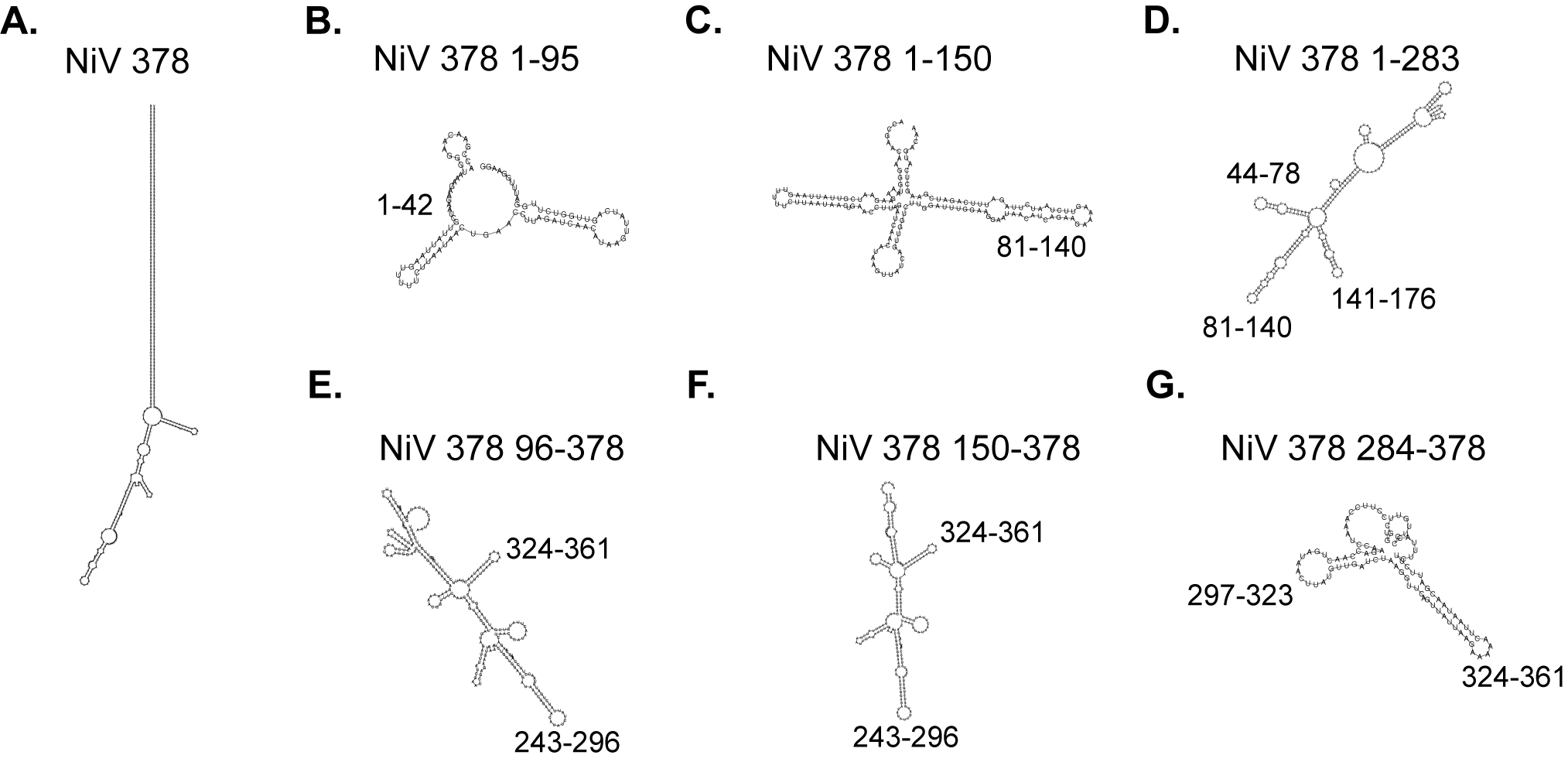

**Figure S4.** **Predicted RNA structures of NiV cbVG 378.** RNAfold predictions for the (A) full-length NiV cbVG 378, (B) nucleotides 1-95 of NiV 378, (C) nucleotides 1-150 of NiV 378, (D) nucleotides 1-283 of NiV 378, (E) nucleotides 96-378 of NiV 378, (F) nucleotides 150-378 of NiV 378, and (G) nucleotides 284-378 of NiV 378. Stem loops found in Fig 3A are indicated.

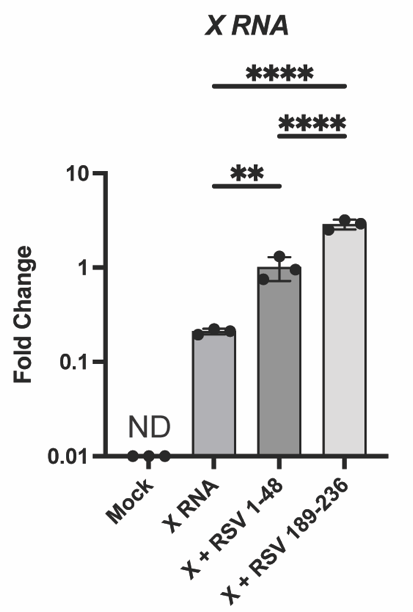

**Figure S5. Transfection efficiency of X RNA constructs.** IVT RNA was transfected into A549 cells and at 6 hpt, RNA was collected and qPCR performed for the X RNA. Data was normalized to the house keeping index of GAPDH and ACTB. One-way ANOVA with Tukey’s HSD posthoc test was performed for statistical analysis. ND: Data was below detection threshold; **: p< 0.01; ****: p<0.0001

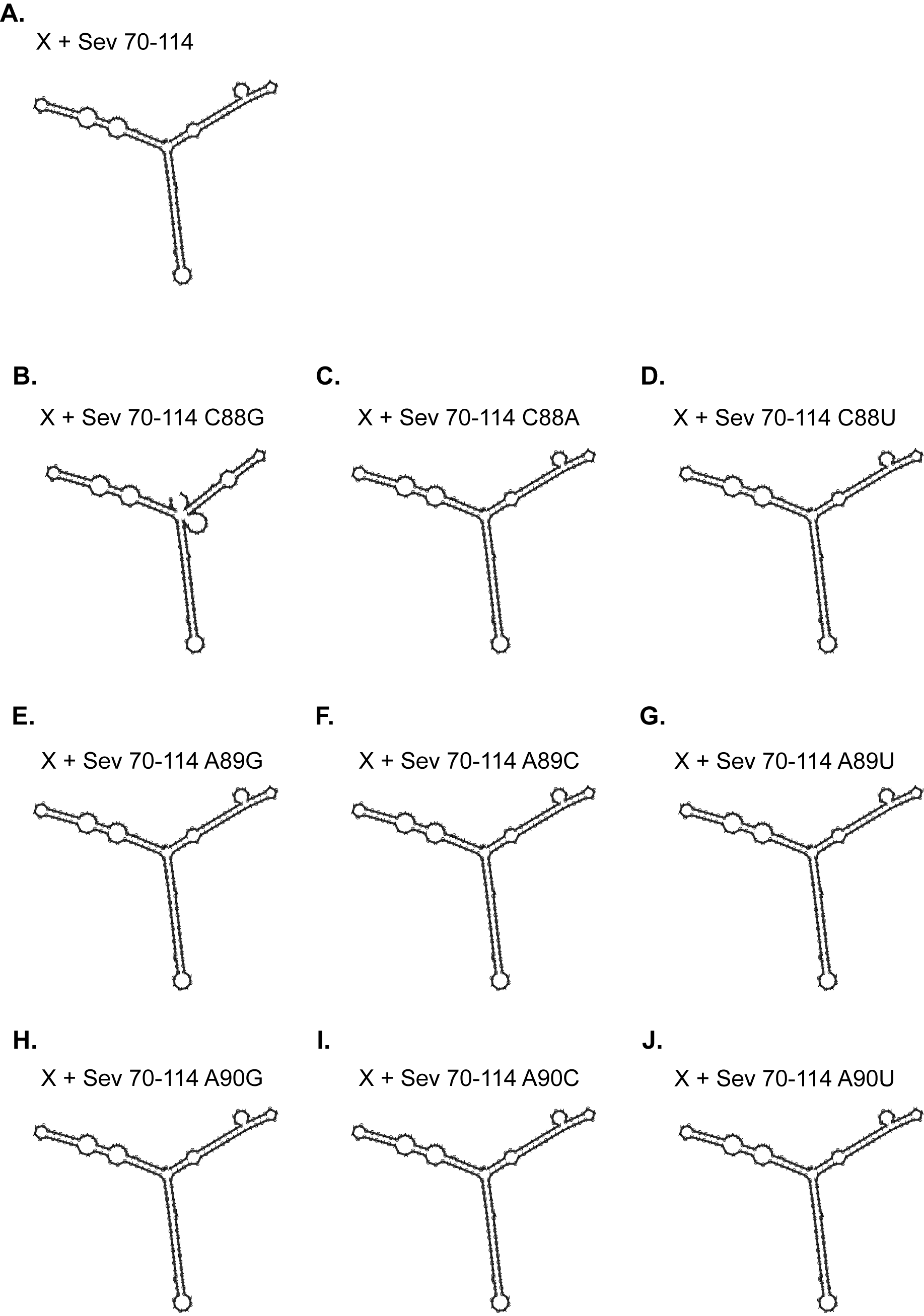

**Figure S6. RNA structures of mutated X RNA-attached SeV 70-114.** RNAfold predictions for (A) X RNA attached with SeV 70-114 stem loop. RNAfold predictions for the X + SeV 70-114 mutations of (B) C88G, (C) C88A, (D) C88U, (E) A89G, (F) A89C, (G) A89U, (H) A90G, (I) A90C, and (J) A90U.

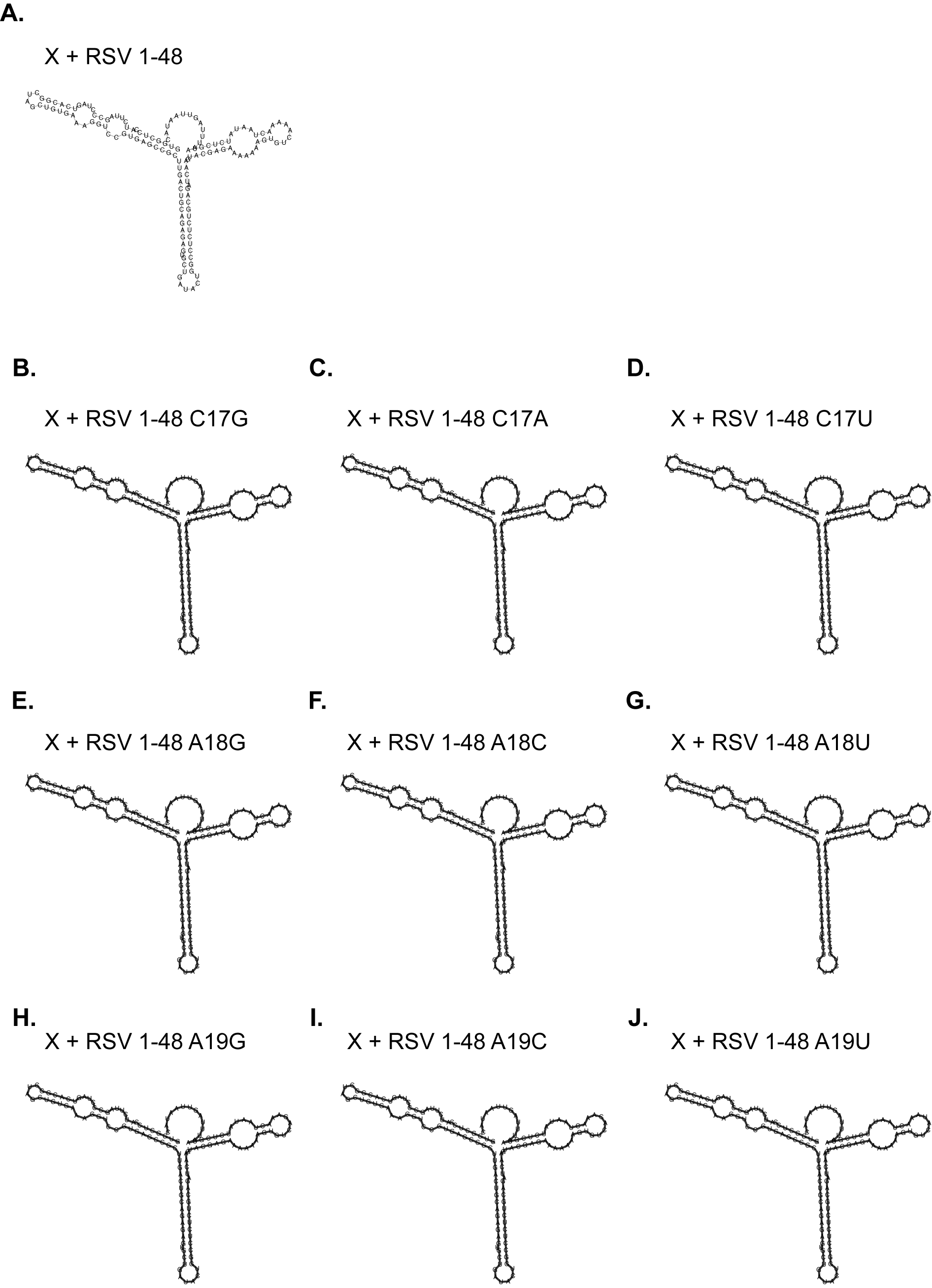

**Figure S7. RNA structures of mutated X RNA-attached RSV 1-34.** RNAfold predictions for (A) X RNA attached with RSV 1-34 stem loop. RNAfold predictions for the X + RSV 1-34 mutations of (B) C17G, (C) C17A, (D) C17U, (E) A18G, (F) A18C, (G) A18U, (H) A19G, (I) A19C, and (J) A19U.

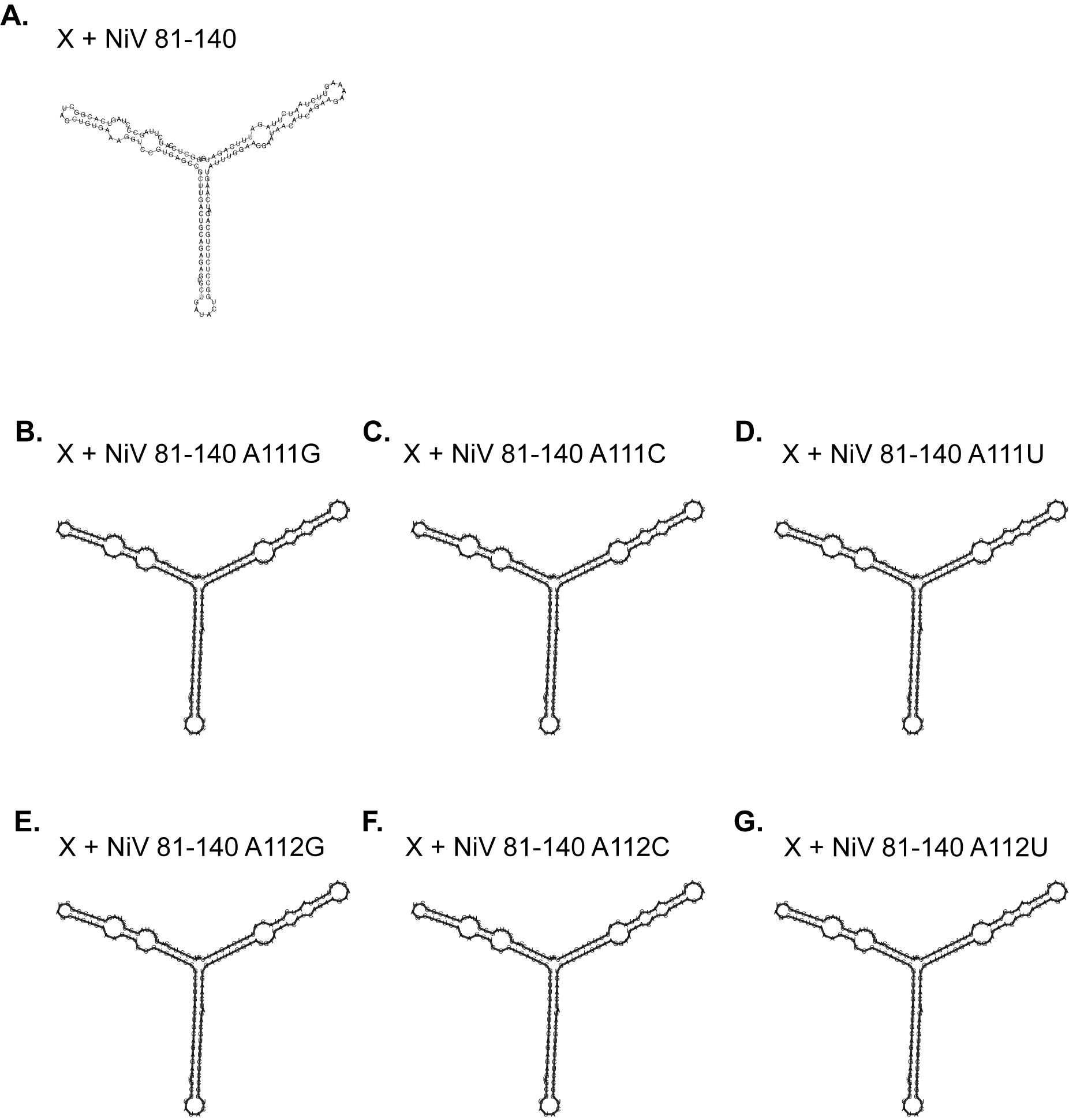

**Figure S8. RNA structures of mutated X RNA-attached NiV 81-140.** RNAfold predictions for (A) X RNA attached with RSV 81-140 stem loop. RNAfold predictions for the X + NiV 81-140 mutations of (B) A111G, (C) A111C, (D) A111U, (E) A112G, (F) A112C, and (G) A112U.
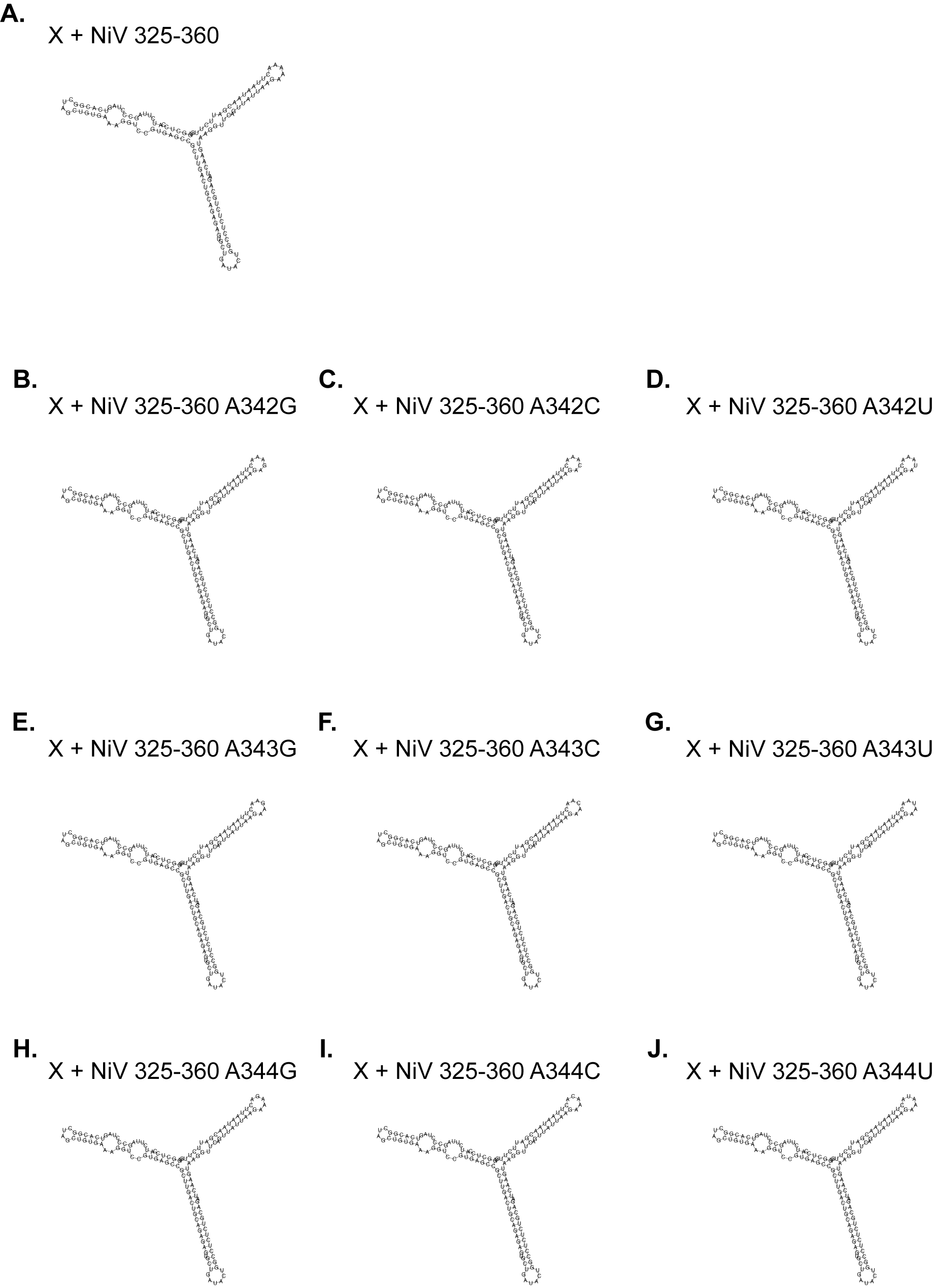

**Figure S9.** **RNA structures of mutated X RNA-attached RSV 325-360.** RNAfold predictions for (A) X RNA attached with RSV 325-360 stem loop. RNAfold predictions for the X + RSV 325-360 mutations of (B) A342G, (C) A342C, (D) A342U, (E) A343G, (F) A343C, (G) A343U, (H) A344G, (I) A344C, and (J) A344U.

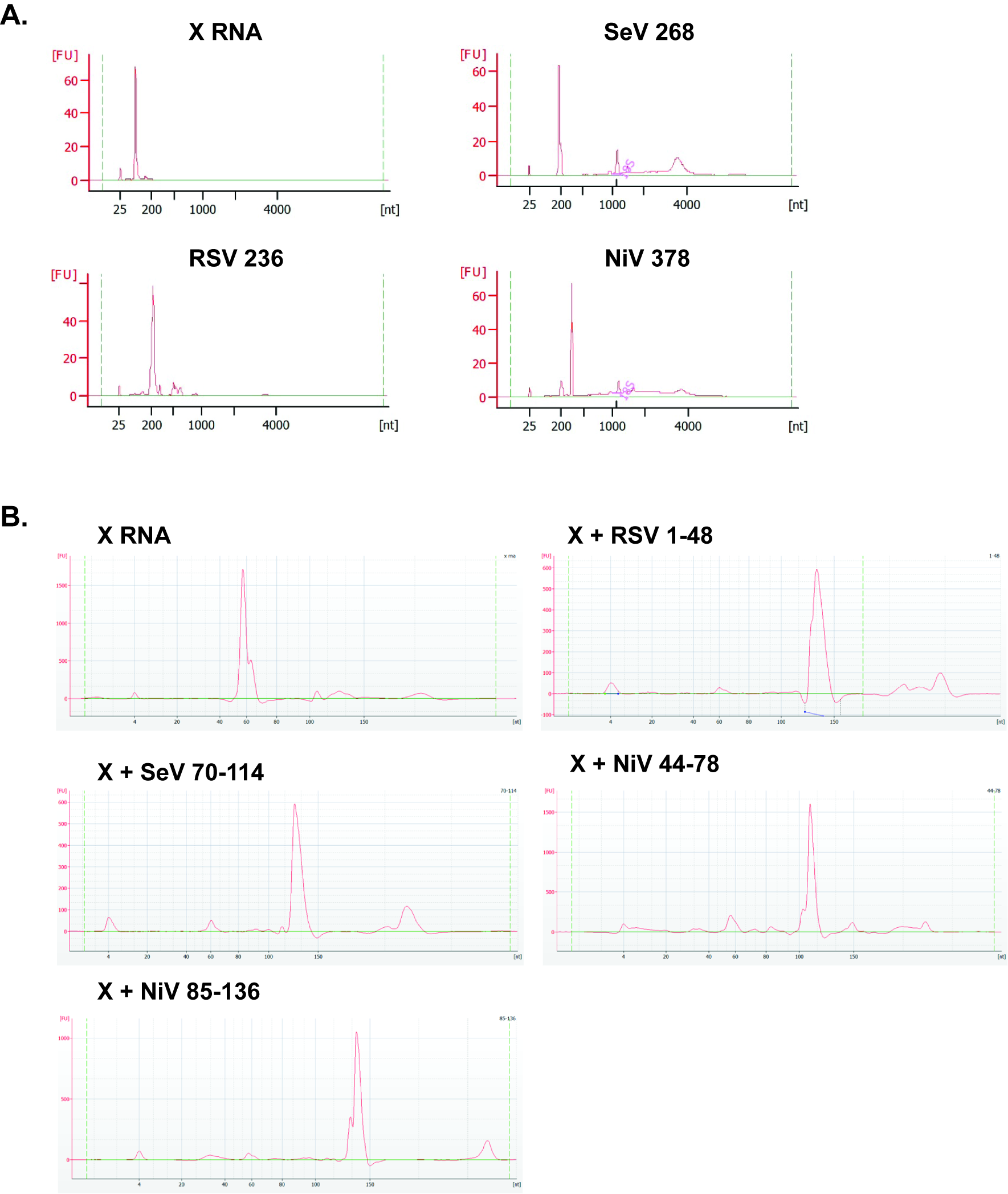

**Figure S10. Bioanalyzer of *In Vitro* Transcribed RNAs**. A) Bioanalyzer curves of cbVGs and the X RNA used in Figure 1. B) Bioanalyzer curves of the X RNA attached constructs.
